# The Alzheimer’s disease associated bacterial protease RgpB from *P. gingivalis* activates the alternative β-secretase meprin β thereby increasing Aβ generation

**DOI:** 10.1101/748814

**Authors:** Fred Armbrust, Cynthia Colmorgen, Claus U. Pietrzik, Christoph Becker-Pauly

## Abstract

Alzheimer’s disease (AD) is the most common type of dementia and characterized by tau hyperphosphorylation, oxidative stress, reactive microglia and amyloid-β (Aβ) deposits. A recent study revealed that *Porphyromonas gingivalis* infection is associated with amyloid β generation in Alzheimer’s disease. Increased Aβ levels, tau degradation and neuronal toxicity were observed as a consequence of ginigipain R (RgpB) activity, a cysteine protease constitutively secreted by *P. gingivalis*. Of note, we previously identified RgpB as a potent activator of the metalloproteinase meprin β. Interestingly, meprin β is an alternative β-secretase of the amyloid precursor protein (APP), which together with the γ-secretase leads to the generation of aggregation-prone N-terminally truncated Aβ2-x peptides. Importantly, identification of a risk gene variant of meprin β (rs173032) for Alzheimer’s disease using whole-exome sequencing of the BDR cohort further supports the impact of this alternative β-secretase. Thus, we wondered if increased Aβ levels as a consequence of *P. gingivalis* colonization into the brain might be due to meprin β activation by RgpB. Here, we demonstrate that i) upon incubation with RgpB the proteolytic activity of meprin β at the cell surface of transfected HEK cells or of endogenously expressed enzyme in SH-SY5Y neuroblastoma cells was significantly increased, and that ii) RgpB-mediated increase in meprin β activity leads to massive generation of Aβ-peptides. In conclusion, our findings would further explain the pathogenesis of *P. gingivalis* in AD brain.

## Background

*Porphyromonas gingivalis* is the major pathogenic bacterium responsible for chronic periodontitis. Potempa and colleagues recently demonstrated that *P. gingivalis* can be found in the brains of Alzheimer’s disease (AD) patients, where it releases the neurotoxic cysteine protease gingipain R (RgpB) [1]. The authors could show that this protease is associated with increased Aβ generation and aggregation, tau degradation and neuronal toxicity.

Despite the exciting results from this paper, we propose an additional molecular mechanism mediated by RgpB being relevant for the neurotoxic activity of *P. gingivalis* infection.

We have demonstrated that RgpB is a highly potent activator of the metalloprotease meprin β at the plasma membrane [2]. Interestingly, meprin β was identified as an alternative β-secretase of the amyloid precursor protein (APP), which together with the γ-secretase leads to the generation of N-terminally truncated Aβ2-x peptides [3,4]. This strongly aggregating Aβ species is particularly increased in AD [5,6], as is meprin β [4,7,8]. Intrigued by the recent study by Potempa and colleagues, demonstrating that RgpB is present in brain tissue of AD patients and associated with pathology [1], we speculated about its effect on meprin β-mediated amyloid β generation.

## Main text

In order to analyze the role of meprin β in Rgpb-induced Aβ generation, we transfected HEK293 cells with APP and/or meprin β and analyzed enzyme activity and APP processing in the presence or absence of RgpB (Fig.1A). Indeed, the amount of Aβ was dramatically increased in the presence of meprin β and the bacterial protease (Fig. 1B). Additionally, meprin β specific generation of a N-terminal APP fragment was also increased upon RgpB treatment (Fig. 1C). Of note, RgpB itself showed proteolytic activity towards APP resulting in degradation of soluble APP ectodomains generated by α- and β-secretases (Fig. 1C). Increased proteolytic activity of meprin β at the cell surface upon RgpB incubation could be measured using a specific peptide cleavage assay [9], further demonstrating the specific interaction between host and pathogenic protease (Fig. 1D). To prove that the enzymatic activity of meprin β was responsible for increased Aβ generation, we applied the meprin inhibitor actinonin [10], which led to strongly decreased Aβ generation in the absence or presence of RgpB (Fig. 1E).

**FIGURE 1.**
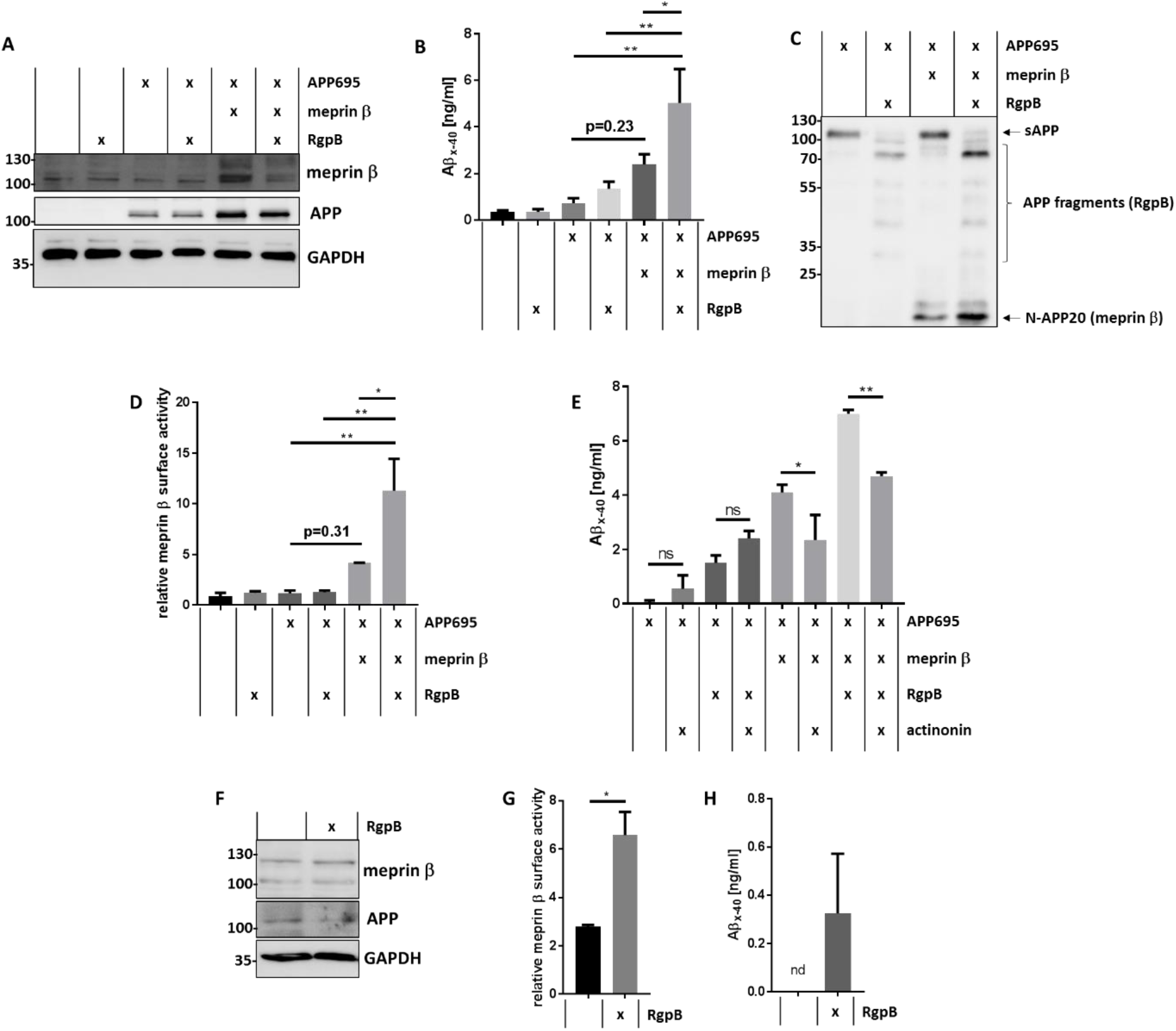
Meprin β activation by RgpB leads to increased Aβ levels. A) HEK cells were transfected with human APP695 and/or human meprin β. After 24 h, cells were treated with 50 nM RgpB for 30 min. Then, the medium was changed to serum-free DMEM. Cells were harvested 3 h later and lysed in an EDTA-containing Triton X-100-based lysis buffer. Proteins were analyzed by Western blot using specific antibodies against meprin β (polyclonal antibody against the ectodomain), APP (CT15, ployclonal antibody against the last C-terminal 15 amino acids of APP) and GAPDH (14C10, Cell Signaling). B) HEK cells were treated as described in A), however, selective samples were treated with E-64 (RgpB inhibitor). Supernatants were analyzed with LEGEND MAX™ β-Amyloid x-40 ELISA Kit (Biolegend) according to manufacturer’s instructions. C) HEK cells were treated as described in A) and supernatants were analyzed by Western blot using an APP antibody (22C11, Biolegend). D) HEK cells were treated as described in A) and proteolytic activity of meprin β at the cell surface was measured using a highly specific quenched fluorogenic peptide substrate ((mca)-EDEDED-(K-e-dnp); mca: 7-methyloxycoumarin-4-yl, dnp: dinitrophenyl) [9]. E) HEK cells were treated as described in A), however, after changing medium to serum-free DMEM, 20 µM of the meprin β inhibitor actinonin was added to selected samples and β-Amyloid x-40 was measured as described in C). F) SHSY5Y cells were treated with 50 nM RgpB for 30 min and meprin β, APP and GAPDH levels were analyzed by Western blot. G) SHSY5Y cells were treated as described in F) and then analyzed for specific meprin β activity as described in D). H) SHSY5Y cells were treated as described in F) and β-Amyloid x-40 was measured as described in C). nd – below detection limit of the ELISA. All statistical analyses were performed with GraphPad Prism using a one-way ANOVA followed by a Turkey’s test for B), D) and E) and a two-tailed t test for G) (*p < 0.05, **p < 0.01).

To analyze a more physiological situation we choose the human neuroblastoma cell line SH-SY5Y, which endogenously express meprin β and APP (Fig. 1F). Employing our peptide cleavage assay we again detected significantly increased meprin β activity at the cell surface when incubated with RgpB (Fig.1G). Of note, we observed Aβ release by RgpB-treated SH-SY5Y cells, whereas Aβ levels generated by untreated SHSY5Y cells were below the detection limit of the ELISA (Fig. 1H).

AD pathogenesis is highly influenced by pro-inflammatory stimuli and subsequent activation of microglia [11]. Several papers proposed bacterial and viral pathogens, such as *Herpes simplex* virus [12], to be involved in the onset of AD. In the recent manuscript published in Science Advances the authors detected the pathogenic bacterium *P. gingivalis*, which is predominantly responsible for periodontitis, also in the brain [1]. In a mouse model, the authors demonstrated that oral infection by *P. gingivalis* indeed leads to induced Aβ peptide formation and destabilization of tau in the murine brain, symptoms that could be diminished applying specific inhibitors of the virulence factors. We provide evidence that *P. gingivalis* induced amyloid plaque formation in AD brain may additionally be mediated through activation of the alternative β-secretase meprin β.

Besides the detrimental role of Aβ and considering meprin β as host-protective enzyme with regard to mucus homeostasis [13,2] and antimicrobial activity [14], one can assume that brain infection by *P. gingivalis* leads to increased activation of meprin β by RgpB and an acute Aβ-generation, which in turn may serve as an antimicrobial peptide [15] (Fig. 2). Of course, there are many open questions that need to be answered and mouse work in this direction is ongoing. However, this is another small piece in the AD puzzle that contributes to our understanding of this devastating disease.

**FIGURE 2.**
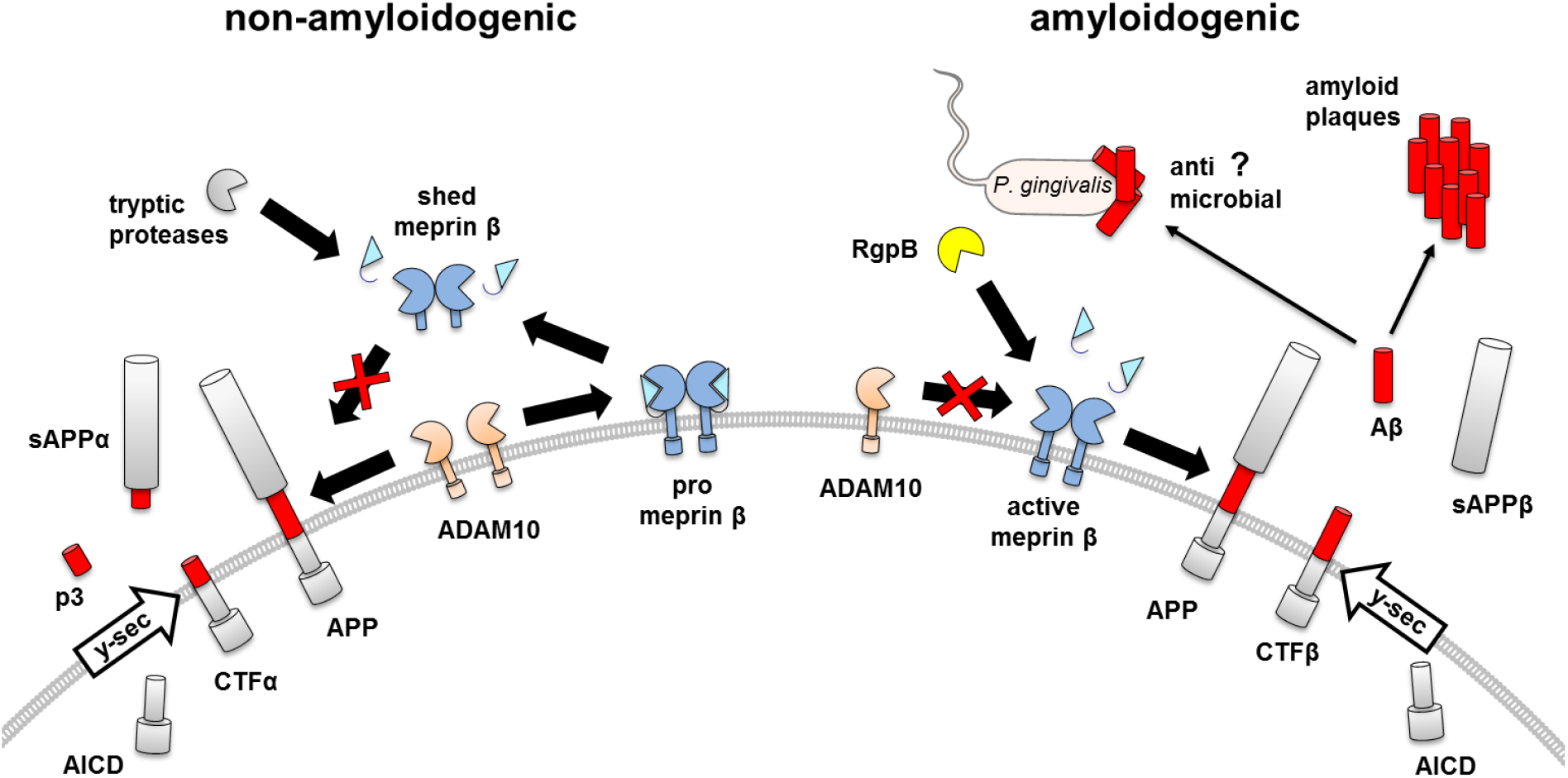
*P. gingivalis* infection promotes the amyloidogenic pathway of APP processing. The alternative β-secretase meprin β can be activated by the pathogenic protease RgpB, secreted from *P. gingivalis*. Activation of meprin β at the cell surface induces Aβ generation. Importantly, only the proform of meprin β is shed by the α-secretase ADAM10, which blocks its β-secretase activity. Hence, activation of meprin β by RgpB has a dual detrimental role, activating the β-secretase and preventing its shedding. However, it is also under debate whether Aβ-peptides might act as anti-microbial peptides in acute bacterial infection.

## Conclusions

Potempa and colleagues observed increased Aβ levels as a consequence of RgpB secretion by *P. gingivalis* being capable of colonizing the brain [1]. Here, we demonstrate that meprin β is involved in the generation of increased Aβ levels upon activation by Rgpb. Our findings attribute meprin β a potential role in the previously reported Rgpb-dependent Aβ increase.

## List of abbreviations

RgpB: ginigipain R
Aβ: amyloid β
AD: Alzheimer’s Disease
APP: amyloid precursor protein

## Declarations

### Ethics approval and consent to participate

Not applicable.

### Consent for publication

Not applicable.

### Availability of data and materials

The datasets used and/or analyzed during the current study are available from the corresponding author on reasonable request.

### Authors’ contributions

FA conducted the experiments, acquired and analyzed the data and wrote the manuscript. CC was involved in conducting experiments. CUP provided regents and revised the manuscript. CBP wrote, edited and revised the manuscript and designed the research study. All authors read and approved the final manuscript. All authors read and approved the final manuscript.

## Acknowledgements

We thank Jan Potempa (University of Louisville) for providing purified RgpB protease from *P. gingivalis*.

